# Polyphosphate acts as an architectural regulator of carbon fixation and nucleoid structure in cyanobacteria

**DOI:** 10.64898/2026.04.09.717567

**Authors:** Claire E. Dudley, Daniel J. Foust, David F. Savage, Julie S. Biteen, Anthony G. Vecchiarelli

## Abstract

Polyphosphate (polyP) is a conserved inorganic polymer traditionally viewed as a stress-induced phosphate and energy reserve. In cyanobacteria, however, polyP granules are constitutively present and frequently observed in proximity to carboxysomes, the bacterial microcompartments that mediate CO_2_ fixation. Here we show that polyP functions as a spatially organized regulator of the photosynthetic cytoplasm in *Synechococcus elongatus*. PolyP granules localize to the nucleoid and are periodically arranged along the cell axis, independently of the McdAB carboxysome positioning system. Despite this independence, polyP and carboxysomes associate non-randomly, and this association is enhanced when active carboxysome positioning by the McdAB system is disrupted. Loss of polyP synthesis leads to nucleoid expansion, an increased number of smaller carboxysomes with high mobility, and severe defects in growth under ambient CO_2_. Perturbation of polyP turnover further reveals structural connections to both carboxysomes and thylakoid membranes. Together, these findings identify polyP as an architectural integrator that couples chromosome organization, metabolic compartmentalization, and photosynthetic fitness.

## INTRODUCTION

Polyphosphate (polyP) is an ancient and highly conserved inorganic polymer composed of linear chains of orthophosphate residues linked by high-energy phosphoanhydride bonds. PolyP granules are found across all three domains of life and have been implicated in diverse aspects of bacterial physiology, including virulence, stress survival, metal homeostasis, and regulation of DNA structure^1–10^. In most heterotrophic bacteria, polyP accumulation is strongly induced by environmental stressors such as nutrient deprivation, osmotic shock, or stringent response activation^5,11–13^. Under these conditions, polyP functions as a phosphate reservoir, an energy buffer, and a stress-response effector.

Intriguingly, in several carboxysome-containing autotrophs - including the γ-proteobacterium *Halothiobacillus neapolitanus* and multiple cyanobacterial species - polyP granules have been observed by electron microscopy in cells grown under favorable laboratory conditions^14–17^. Unlike stress-induced polyP accumulation in heterotrophs, these granules appear constitutively present in exponentially growing autotrophic cells. The biological necessity for polyP in these unstressed conditions remains unclear. Surprisingly, ultrastructural studies have reported physical proximity between polyP granules and carboxysomes, sometimes connected by unidentified “lattice-like” structures^16,18^. Whether this association is incidental or functionally meaningful is unknown. Thus, the mechanistic basis and physiological significance of the polyP–carboxysome relationship remain unresolved.

Carboxysomes are proteinaceous bacterial microcompartments that encapsulate Rubisco and Carbonic Anhydrase to enhance CO_2_ fixation efficiency^16^. In the model cyanobacterium *Synechococcus elongatus* PCC 7942 (hereafter *S. elongatus*), carboxysomes are not randomly distributed within the cytoplasm but are actively positioned along the cell axis by the maintenance of carboxysome distribution (Mcd) system^19,20^. This two-protein positioning system consists of the ParA-like ATPase McdA and its cargo-linking partner McdB. In its ATP-bound state, McdA binds nonspecifically to the nucleoid, while McdB associates with carboxysomes and stimulates McdA ATP hydrolysis, generating a self-organizing gradient that evenly distributes carboxysomes^19,20^. We have further shown that nucleoid compaction state directly influences carboxysome localization and dynamics, revealing that the nucleoid acts as a positioning matrix for these metabolic organelles^22^. Whether other regularly organized cellular structures, such as polyP granules, are coordinated with this nucleoid-based positioning system is unknown.

PolyP metabolism is governed primarily by polyphosphate kinases (PPKs) and exopolyphosphatases (PPXs). PPK1 is typically the major enzyme responsible for polyP synthesis, while PPK2 enzymes can contribute to both synthesis and degradation depending on cellular context^21,22^. The number and specialization of PPK enzymes vary widely across bacteria. For example, *Escherichia coli* encodes a single PPK1, whereas *Pseudomonas aeruginosa* possesses one PPK1 and multiple PPK2 paralogs, with partially redundant functions^5,23^. This diversity underscores that although polyP itself is highly conserved, its physiological deployment is evolutionarily flexible. *S. elongatus* encodes a single predicted PPK1, PPK2, and PPX, yet their respective contributions to polyP granule formation, turnover, and cellular physiology remain uncharacterized.

Beyond its canonical role in phosphate storage, emerging evidence suggests that polyP can influence nucleoid architecture^8^. PolyP has been shown to interact with nucleoid-associated proteins and to promote heterochromatin-like DNA compaction in bacteria^7^. Given that carboxysome positioning depends on nucleoid organization^35^, polyP could plausibly influence carboxysomes indirectly by modulating chromosome structure. Additionally, photosynthesis generates large fluctuations in ATP and phosphate flux, potentially linking polyP metabolism to both light-driven energy production and carbon fixation^24–28^. Photosynthetic ATP powers high-affinity phosphate uptake and polyP synthesis during phosphate-replete conditions, whereas polyP hydrolysis during phosphate limitation can release inorganic phosphate (Pi) and potentially buffer energy imbalances^29–32^. Thus, polyP sits at the intersection of phosphate homeostasis, energy metabolism, and nucleoid organization in photoautotrophs.

Here, we investigate the spatial and functional relationship between polyP granules and carboxysomes in *S. elongatus*. We show that polyP granules are nucleoid-associated and periodically organized along the cell axis. While we find that carboxysomes frequently associate with polyP granules *in vivo*, polyP does not require carboxysomes or McdAB-mediated positioning for its organization. In fact, in the absence of active McdAB positioning, carboxysomes cluster more frequently around polyP granules, revealing that the positioning system normally restrains an intrinsic affinity between these structures. Genetic disruption of polyP synthesis via deletion of *ppK1* abolishes polyP granules and results in nucleoid expansion, altered carboxysome organization, and severe growth defects under ambient CO_2_. Perturbation of polyP turnover further reveals structural and physiological effects on both carboxysomes and thylakoid membranes, linking polyP metabolism to both the dark and light reactions of photosynthesis.

Together, our findings uncover a previously unrecognized role for polyP as a spatial and metabolic regulator of the photosynthetic cytoplasm. Rather than functioning solely as a stress-induced storage polymer, polyP emerges as an architectural integrator that couples nucleoid compaction state, carboxysome organization, and photoautotrophic growth. These findings broaden our understanding of bacterial subcellular organization and establish metabolic polymers like polyP as active, global architects of intracellular structure rather than passive storage molecules.

## RESULTS

### PolyP granules localize to the nucleoid in *S. elongatus* cells

In cyanobacteria and other autotrophic bacteria, electron microscopy studies have reported close physical associations between polyP granules and carboxysomes^14–16,18,33^. However, the mechanism underlying this association, and whether polyP positioning and function are somehow spatially coordinated with carboxysomes, remains unclear. Because carboxysomes are dynamically organized within the cytoplasm^19,20,34,35^, determining the spatial distribution of polyP in live cells is a critical first step toward understanding this relationship. We therefore established a robust method to visualize polyP granules in live *S. elongatus* cells.

In *Synechocystis sp.* PCC 6803, DAPI staining has been used to label polyP granules^36^. We adapted and optimized a similar protocol for *S. elongatus* (see Methods). Following four hours of DAPI staining, polyP granules were visible in the DAPI channel together with the nucleoid (**Fig. 1A**). Importantly, when DAPI binds polyP, its emission spectrum undergoes a red-shift, enabling selective visualization of polyP granules in filter sets configured for CFP^37^. Using this spectral distinction, we observed that polyP granules consistently localize over the DAPI-stained nucleoid (**Fig. 1A**). Because *S. elongatus* is polyploid, harboring approximately 2–10 chromosome copies depending on cell cycle stage^38–40^, we considered the possibility that polyP granules localized within DNA-free gaps between chromosomes rather than directly associating with chromosomal DNA. To test this hypothesis, we treated cells with ciprofloxacin, a gyrase inhibitor previously shown to compact the nucleoid in *S. elongatus*^35,41^. Under these conditions, polyP granules remained associated with the condensed nucleoid mass **(Fig. S1A–B)**.

**Figure 1.**
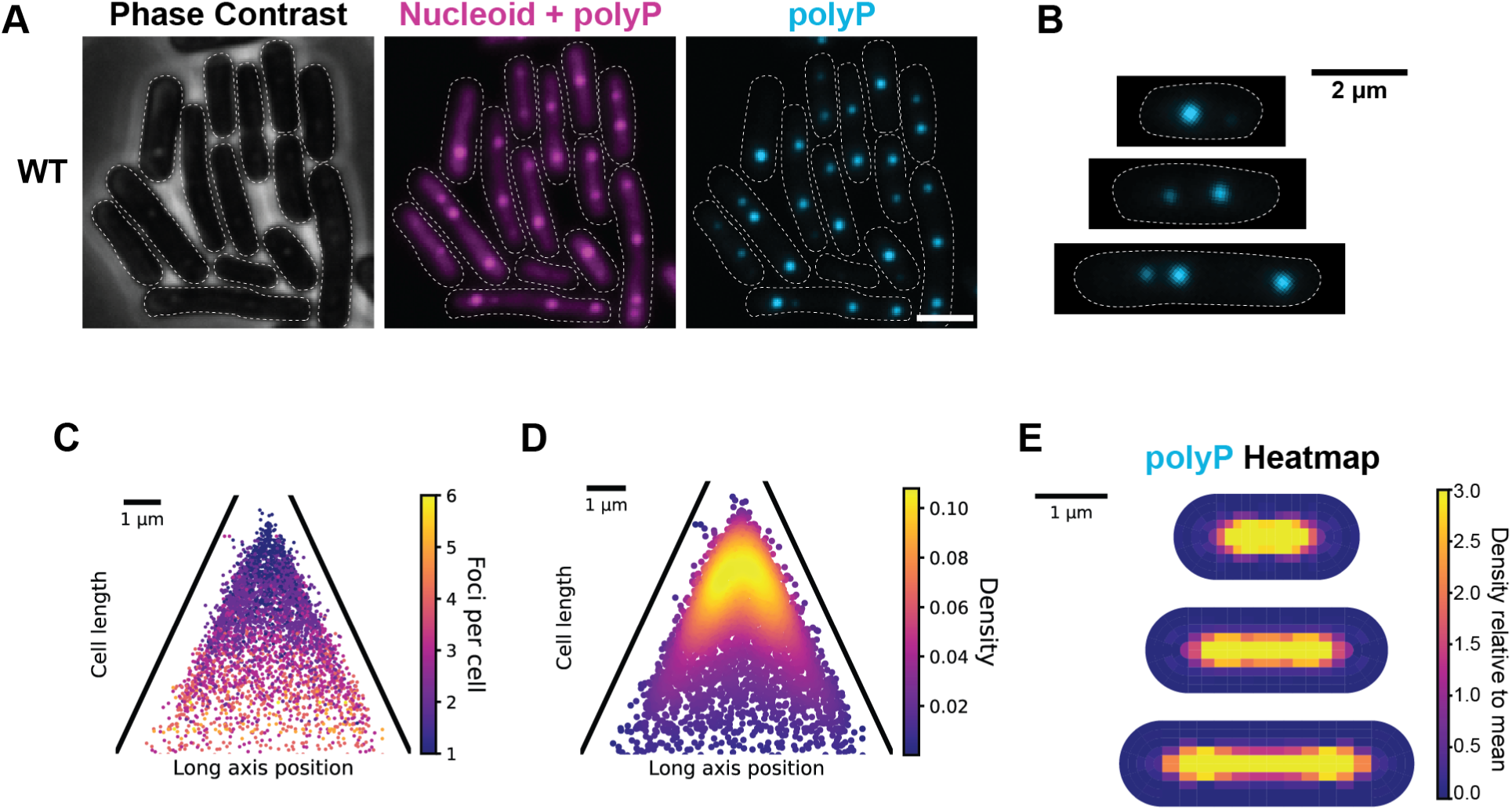
PolyP granules distribute over the nucleoid. **(A)** Representative microscopy images of WT exponential cells (phase contrast), stained with DAPI (magenta), to visualize polyP (cyan), dotted white line represents the cell boundary from phase contrast microscopy (scale bar, 2 μm). **(B)** polyP granules (cyan) in cells of varying cell length, dotted white line represents the cell boundary from phase contrast. **(C)** Number of polyP granules per cell plotted as a function of cell length and long axis position. **(D)** Heat map representing the average localization of polyP granules within WT exponential phase cells. **(E)** polyP heatmap illustrating the probability of localization of polyP granules for cells grouped into quartiles by length from cells analyzed.

Newborn cells typically had one polyP granule localized at midcell, and as cell length increased, so did the number of polyP granules **(Fig. 1B–E)**. In longer cells, two polyP granules were typically positioned near the 1/4 and 3/4 positions along the cell axis **(Fig. 1B)** — a spatial pattern observed in other bacteria^5,12,23^. In longer, pre-divisional cells, as many as 3 to 6 polyP granules were distributed across the cell length **(Fig. 1B-E)**. Together, these results show that polyP granules are spatially organized in close association with the nucleoid of *S. elongatus*.

### PolyP granules associate with carboxysomes

The periodic positioning of polyP granules along the nucleoid suggested that their localization may be actively coordinated with other intracellular structures. Previous electron microscopy studies have reported physical proximity between polyP granules and carboxysomes^14–16,18,33^, but this association has yet to be observed and quantified in living cells. We therefore directly imaged polyP granules and carboxysomes in live *S. elongatus* cells using fluorescence microscopy.

Both RbcS–mOrg carboxysomes and DAPI-stained polyP granules were visualized within the same cells **(Fig. 2A)**. Within individual cells, multiple instances of carboxysomes associating with polyP granules were observed **(Fig. 2A, arrows)**. Across the population, both structures were distributed across the cell axis **(Fig. 2B–E)**. However, there were significantly more carboxysomes than polyP granules in most cells **(Fig. 2F)**. Therefore, only a subset of carboxysomes associated with polyP granules.

**Figure 2.**
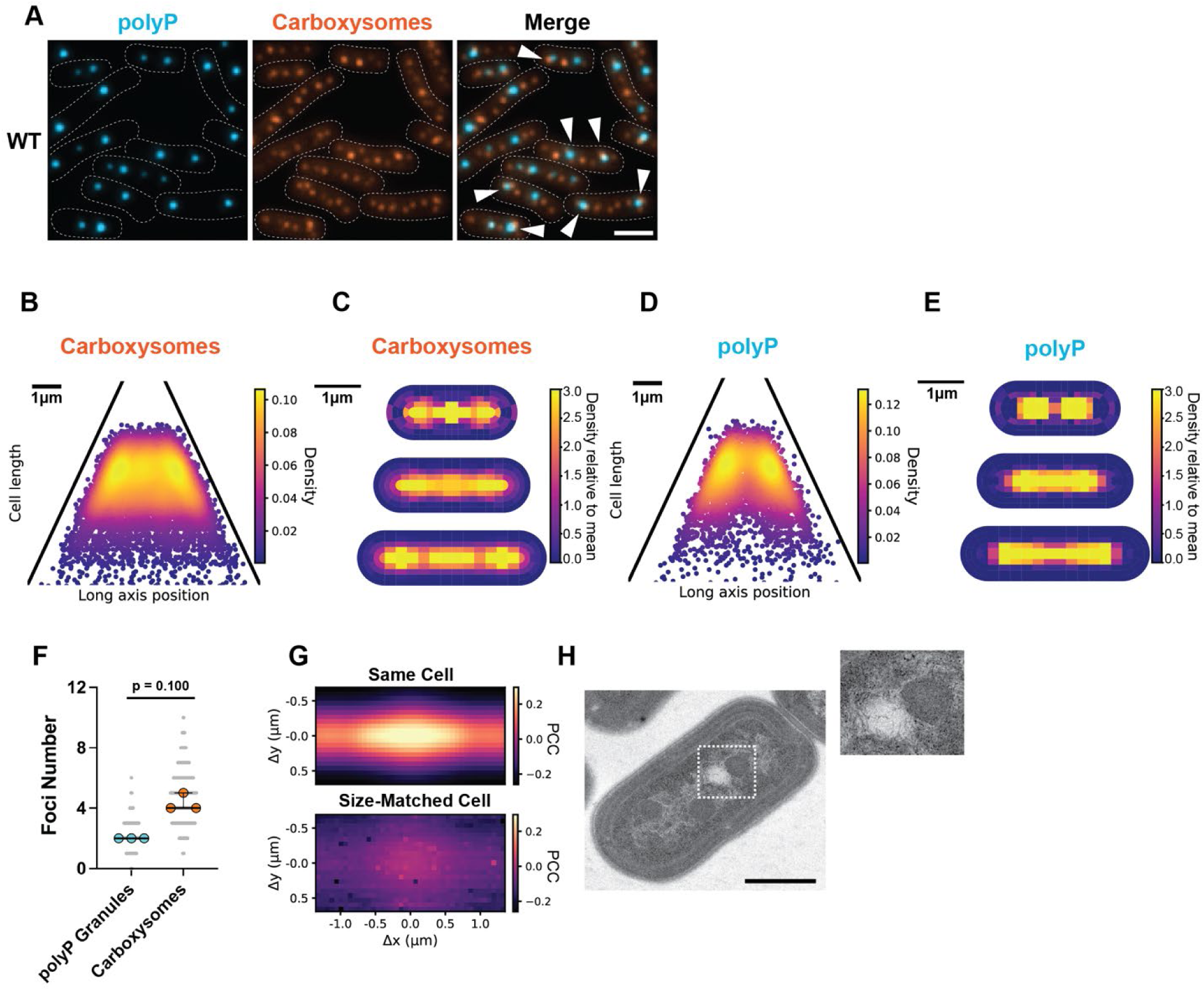
PolyP granules associate with carboxysomes. **(A)** Microscopy images of polyP granules (cyan), carboxysomes (orange), and the merge showing overlap of the two channels. Arrows indicate events of polyP-carboxysome interaction, dotted white line represents the cell boundary from phase contrast (scale bar, 2 μm). **(B)** Heat map representing the average localization of carboxysomes within WT exponential phase cells. **(C)** Probability localization heatmaps of carboxysomes for cells grouped into quartiles by length from cells analyzed. **(D)** Heat map representing the average localization of polyP granules within WT exponential phase cells. **(E)** polyP heatmap illustrating the probability of localization of polyP granules for cells grouped into quartiles by length from cells analyzed. **(F)** Focus number of polyP granules and carboxysomes, Mann-Whitney t-test, p = 0.100, n = 3 biological replicates, bars show median and interquartile range. **(G)** Normalized cross-correlation analysis (see description in Methods section) of carboxysomes and polyP granules from the same cell (top panel) and size-matched cell (bottom panel). Color indicates the Pearson correlation coefficient (PCC), and the x- and y-axes demonstrate shifts of the carboxysome channel. **(H)** Transmission electron micrograph of WT exponential *S. elongatus* cell. Inset illustrating interaction between carboxysome and polyP granule (scale bar, 500 nm).

To quantitatively assess their spatial relationship, we performed normalized cross-correlation analysis. For each cell, the Pearson correlation coefficient (PCC) was calculated between the carboxysome and polyP imaging channels; PCC values approaching 1 indicate strong colocalization. We then computationally shifted the carboxysome channel in single-pixel increments along the x- and y-axes and recalculated the PCC at each displacement. This analysis generated a broad correlation peak centered near zero shift **(Fig. 2G)**, indicating that polyP granules are indeed frequently located in close proximity to carboxysomes. The width of the peak is consistent with adjacent positioning rather than precise one-to-one overlap. To determine whether this correlation could arise from random intracellular positioning, we performed a control analysis in which the polyP channel from a different cell of comparable size (‘size-matched cell’) was paired with the carboxysome channel. Under these conditions, the correlation was lost and PCC values were markedly reduced **(Fig. 2G)**, demonstrating that the observed association is significant and not attributable to random spatial overlap.

To further examine the polyP-carboxysome association at higher resolution, we performed transmission electron microscopy on wildtype (WT) exponentially growing cells. As shown previously^14,20^, carboxysomes were distributed along the cell length, and instances of carboxysomes physically associating with polyP granules were observed **(Fig. 2H)**, consistent with our fluorescence microscopy findings.

Collectively, these data demonstrate that polyP granules and carboxysomes spatially associate in living *S. elongatus* cells at a frequency exceeding that expected from independent intracellular positioning. This non-random organization supports a model in which polyP granule and carboxysome organization is coordinated within the cytoplasm and suggests a functional or mechanistic link between phosphate storage and carbon fixation.

### PolyP organization does not require carboxysomes or the McdAB system

The periodic organization of polyP granules along the nucleoid closely resembles the spacing pattern of carboxysomes, which are actively positioned on the nucleoid by the McdAB system. We therefore hypothesized that polyP granules might associate with carboxysomes to achieve their regular spacing by hitchhiking on their McdAB-dependent positioning mechanism. We first determined whether the McdAB positioning system influences polyP localization. In Δ*mcdA*, Δ*mcdB*, and Δ*mcdAB* backgrounds, carboxysomes cluster into mispositioned aggregates^20,34^, but polyP granules remained normally organized, with no significant changes in number, intensity, or overall spatial distribution relative to WT cells **(Fig. 3A-D, F-I)**. As shown previously, Δ*mcdA* cells are shorter^20,34^, but this defect in cell morphology does not disrupt polyP organization in the cell **(Fig. 3F-I)**.

**Figure 3.**
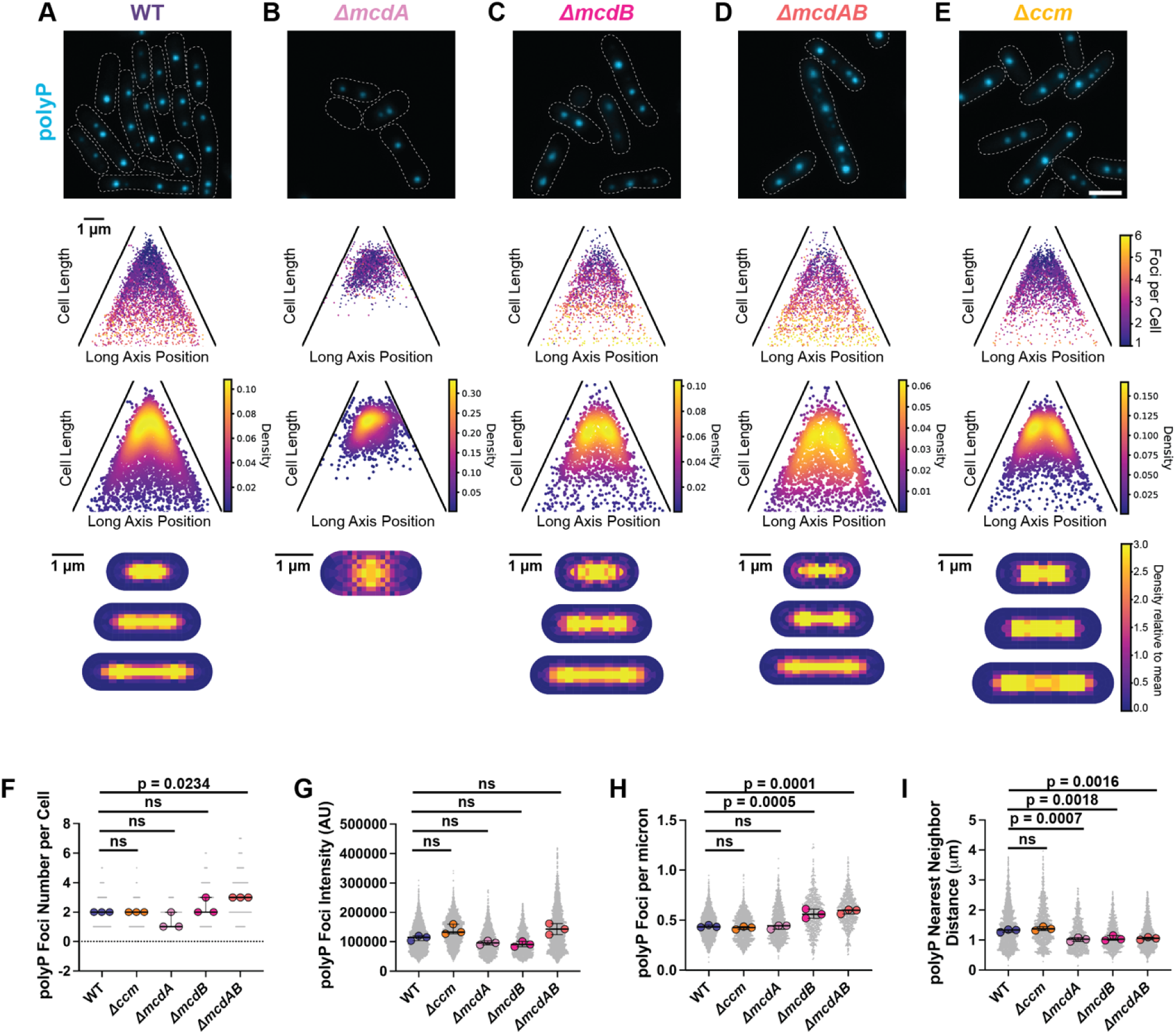
PolyP organization is not dependent on carboxysomes. (A-E) Microscopy images of polyP granules (cyan) in WT, Δ*mcdA*, Δ*mcdB*, Δ*mcdAB*, and Δ*ccm* (carbon concentrating mechanism), dotted white line represents the cell boundary from phase contrast (scale bar, 2 μm). (A) is the same microscopy image and plots from Figure 1ACDE. Number of polyP granules per cell plotted as a function of cell length and long axis position. Heat map representing the average localization of polyP granules. PolyP heatmap illustrating localization probability of polyP granules for cells grouped into quartiles by length from cells analyzed. **(F)** Number of polyP granules per cell. Significance from one-way ANOVA and Dunnett’s multiple comparisons. n = 3 biological replicates, bars show median and interquartile range. **(G)** Intensity of polyP foci (arbitrary units). Significance from one-way ANOVA and Dunnett’s multiple comparisons. n = 3 biological replicates, bars show median and interquartile range. **(H)** PolyP granules per unit of cell length, calculated by dividing the number of foci by cell length. Significance from one-way ANOVA and Dunnett’s multiple comparisons. n = 3 biological replicates, bars show median and interquartile range. **(I)** Distance between polyP granule and its nearest polyP neighbor. Significance from one-way ANOVA and Dunnett’s multiple comparisons. n = 3 biological replicates, bars show median and interquartile range.

We next examined whether carboxysomes themselves are required for polyP organization by imaging polyP granules in an *S. elongatus* strain devoid of carboxysomes. The Δ*ccm* strain lacks the shell proteins essential for carboxysome assembly^20,42^. In the absence of carboxysomes, polyP granules remained regularly positioned over the nucleoid along the cell length with spacing, copy number, and fluorescence intensities comparable to those of WT cells **(Fig. 3A and E, F-I)**. Together, these findings demonstrate that, despite their nucleoid-associated distribution and physical association with carboxysomes, polyP granules do not depend on carboxysomes or McdAB-mediated positioning for their formation or spatial organization. Instead, polyP distribution is governed by a distinct, yet-to-be discovered, nucleoid-dependent mechanism independent of the McdAB carboxysome positioning system.

### Carboxysomes cluster around polyP granules in the absence of the McdAB system

Having established that carboxysome assembly and positioning have no major influence on polyP organization, we next asked whether the spatial relationship between carboxysomes and polyP changes when carboxysomes are mispositioned. To simultaneously image both structures, we generated fluorescently labeled carboxysomes (RbcS-mOrg) in McdAB system mutant strains. As shown previously^20,34^, carboxysomes form mispositioned clusters in Δ*mcdA*, Δ*mcdB*, or Δ*mcdAB* cells **(Fig. 4A)**. Strikingly, these aggregated carboxysomes associated more frequently with polyP granules than in WT cells **(Fig. 4A)**, as indicated by the significantly narrower cross-correlation peaks for the mutant strains **(Fig. 4B)**.

**Figure 4.**
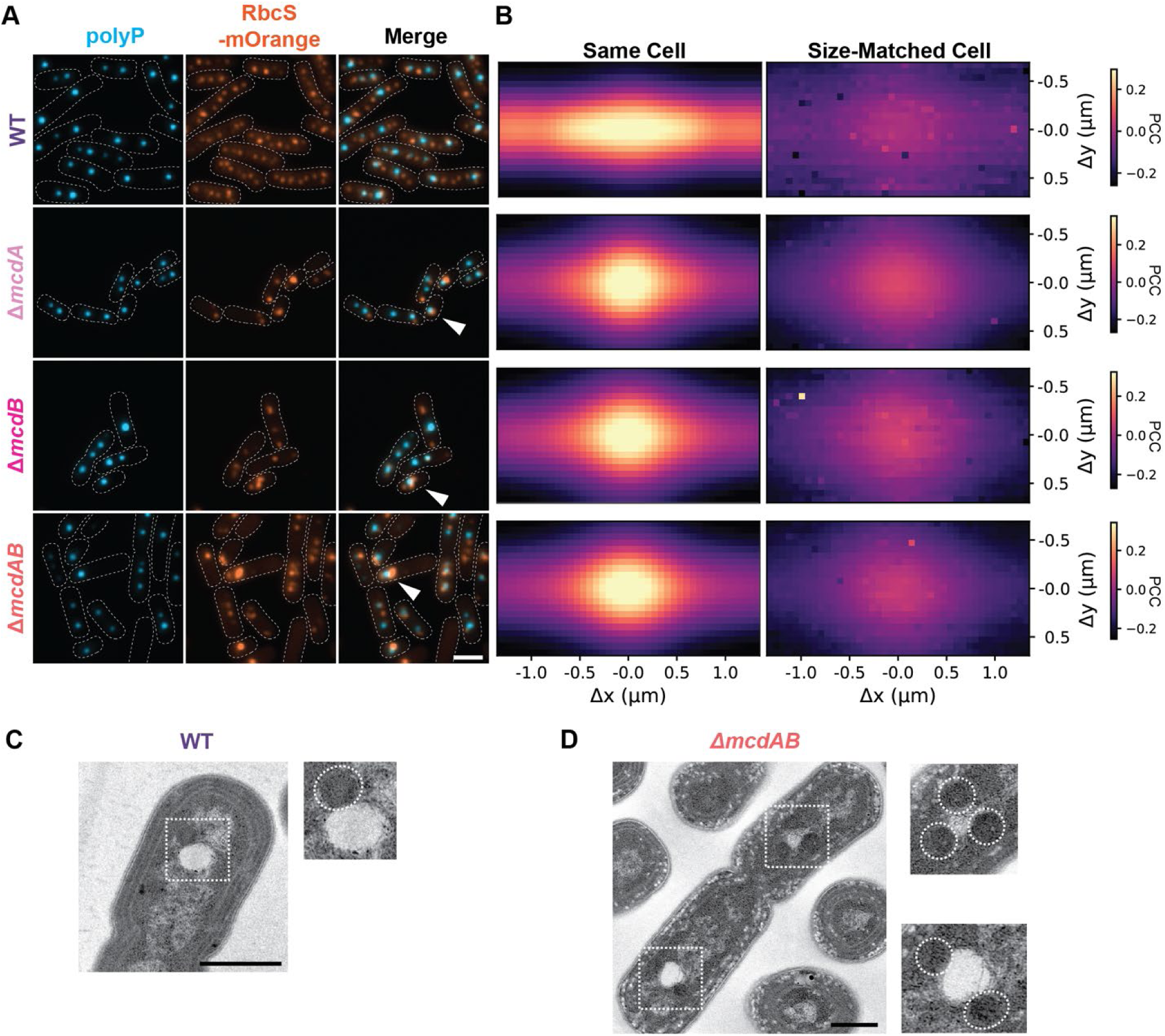
Carboxysomes cluster around polyP granules in the absence of active positioning by the McdAB system. **(A)** Microscopy images of polyP granules (cyan), carboxysomes (orange), and the merge shows the overlap of the 2 channels. Arrows indicate events of polyP-carboxysome interaction, dotted white line represents the cell boundary from phase contrast (scale bar, 2 μm). WT images are the same as Figure 2A. **(B)** Normalized cross-correlation analysis (see description in Methods section) of carboxysomes and polyP granules from the same cell (top panel) and size-matched cell (bottom panel) for WT, Δ*mcdA*, Δ*mcdB*, Δ*mcdAB*. Color indicates the Pearson correlation coefficient (PCC) and the x- and y- axes demonstrate shifts of the carboxysome channel. WT panels are duplicates of Figure 2G. **(C)** Transmission electron micrograph of WT exponential phase cell. Inset illustrates carboxysome-polyP interaction (scale bar, 600 nm). **(D)** Transmission electron micrograph of Δ*mcdAB* exponential phase cell. Insets illustrate carboxysome-polyP interactions. White dashed circles indicate carboxysomes. Scale bar: 500 nm.

To spatially resolve this interaction beyond the limits of fluorescence microscopy, we performed transmission electron microscopy. In exponentially growing WT cells, carboxysome–polyP contacts were infrequent and discrete **(Fig. 4C)**, whereas Δ*mcdAB* cells displayed multiple carboxysomes clustered around single polyP granules, corroborating the fluorescence analysis **(Fig. 4D)**. Collectively, these results indicate that, in addition to distributing carboxysomes across the nucleoid, the McdAB system prevents carboxysome coalescence at polyP granules.

### Drug-induced nucleoid compaction reduces carboxysome clustering at polyP granules

We previously demonstrated that, in the absence of McdAB-mediated positioning, carboxysomes form nucleoid-excluded aggregates — a phenotype exacerbated by ciprofloxacin-induced nucleoid compaction^41^. To further examine the spatial relationships among carboxysomes, polyP granules, and the nucleoid, we treated WT and Δ*mcdAB* cells with ciprofloxacin. In WT cells, both carboxysomes and polyP granules remained associated with the nucleoid following treatment **(Fig. S2A-B)**. In contrast, in Δ*mcdAB* cells, some carboxysome foci became nucleoid-excluded while polyP granules remained tightly associated with the hyper-compacted nucleoid. Together, these observations indicate that polyP–nucleoid interactions are more robust than polyP–carboxysome associations, reinforcing a model in which nucleoid coupling is the primary determinant of polyP spatial organization.

### PPK1 is required for polyP granule formation in *S. elongatus*

We next genetically disrupted polyP metabolism to define the pathway responsible for granule formation and to establish conditions lacking polyP for downstream analyses on carboxysome organization and function. To this end, we generated deletion mutants targeting enzymes implicated in polyP synthesis and degradation. The native loci encoding the polyphosphate kinases PPK1 and PPK2, and the exopolyphosphatase PPX (SYNPCC7942_1566, SYNPCC7942_0493, and SYNPCC7942_1965), were replaced with antibiotic resistance cassettes to create Δ*ppK1*, Δ*ppK2*, and Δ*ppX* strains. Because PPK2 can contribute to polyP turnover under certain conditions^21,22^, we additionally constructed a Δ*ppK2*Δ*ppX* double mutant to further restrict polyP degradation.

All mutants except Δ*ppK1* retained detectable polyP granules **(Fig. 5A-B)**. When present, polyP distribution and copy number were comparable to WT **(Fig. 5C–E)**. Notably, polyP granules in the Δ*ppK2*Δ*ppX* mutant were brighter than in WT **(Fig. 5F)**, consistent with impaired polyP degradation. In contrast, Δ*ppK1* cells lacked detectable polyP granules by both electron microscopy and DAPI fluorescence imaging **(Fig. 5A-B and E)**, identifying PPK1 as the principal driver of polyP synthesis in *S. elongatus*. Notably, Δ*ppK1* cells also displayed increased cell length, width, and area relative to WT **(Fig. S3A–C)**, which intriguingly are also hallmark carbon-limitation phenotypes we previously observed in *mcd* mutants with aggregated and mispositioned carboxysomes^34^.

**Figure 5.**
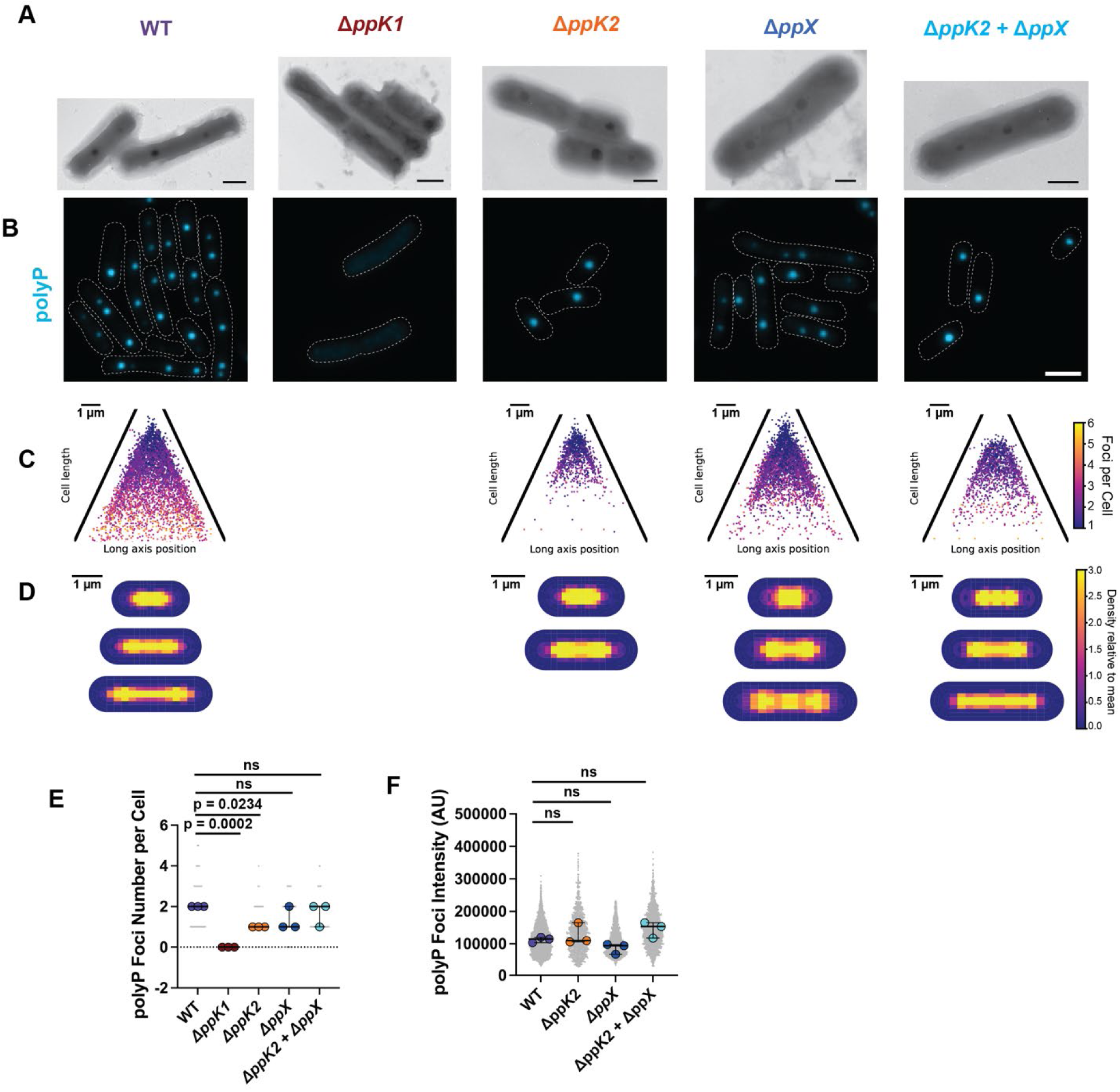
PPK1 perturbations disrupts polyP formation. **(A)** Whole cell electron micrographs of WT, Δ*ppK1*, Δ*ppK2*, Δ*ppX*, and Δ*ppK2*Δ*ppX* without fixative (see methods for preparation) (scale bars, WT - 600 nm, Δ*ppK1* - 1 μm, Δ*ppK2* - 600 nm, Δ*ppX* - 400 nm, and Δ*ppK2*Δ*ppX* - 600 nm). **(B)** Microscopy of polyP granules (cyan) in WT, Δ*ppK1*, Δ*ppK2*, Δ*ppX*, and Δ*ppK2*Δ*ppX* cells, dotted white line represents the cell boundary from phase contrast (scale bar, 2 μm). WT images are the same as Figure 1A. **(C)** Number of polyP granules per cell plotted as a function of cell length and long axis position for WT, Δ*ppK1*, Δ*ppK2*, Δ*ppX*, and Δ*ppK2*Δ*ppX* cells. WT plot is the same as Figure 1C. **(D)** PolyP heatmaps illustrating the localization probability of polyP granules for cells grouped into quartiles by length from cells analyzed for WT, Δ*ppK1*, Δ*ppK2*, Δ*ppX*, and Δ*ppK2*Δ*ppX* cells. WT plot is the same as Figure 1E. **(E)** Number of polyP granules per cell. Significance from one-way ANOVA and Dunnett’s multiple comparisons. n = 3 biological replicates, bars show median and interquartile range. **(F)** Intensity of polyP foci (arbitrary units). Significance from one-way ANOVA and Dunnett’s multiple comparisons. n = 3 biological replicates, bars show median and interquartile range.

### Loss of PPK1 disrupts carboxysome organization and nucleoid compaction

Having established that Δ*ppK1* eliminates polyP granules, we next examined if the loss of polyP affects carboxysome organization. PolyP mutants were introduced into an RbcS-mOrg background to simultaneously visualize DAPI-stained polyP and carboxysomes **(Fig. 6A)**.

**Figure 6.**
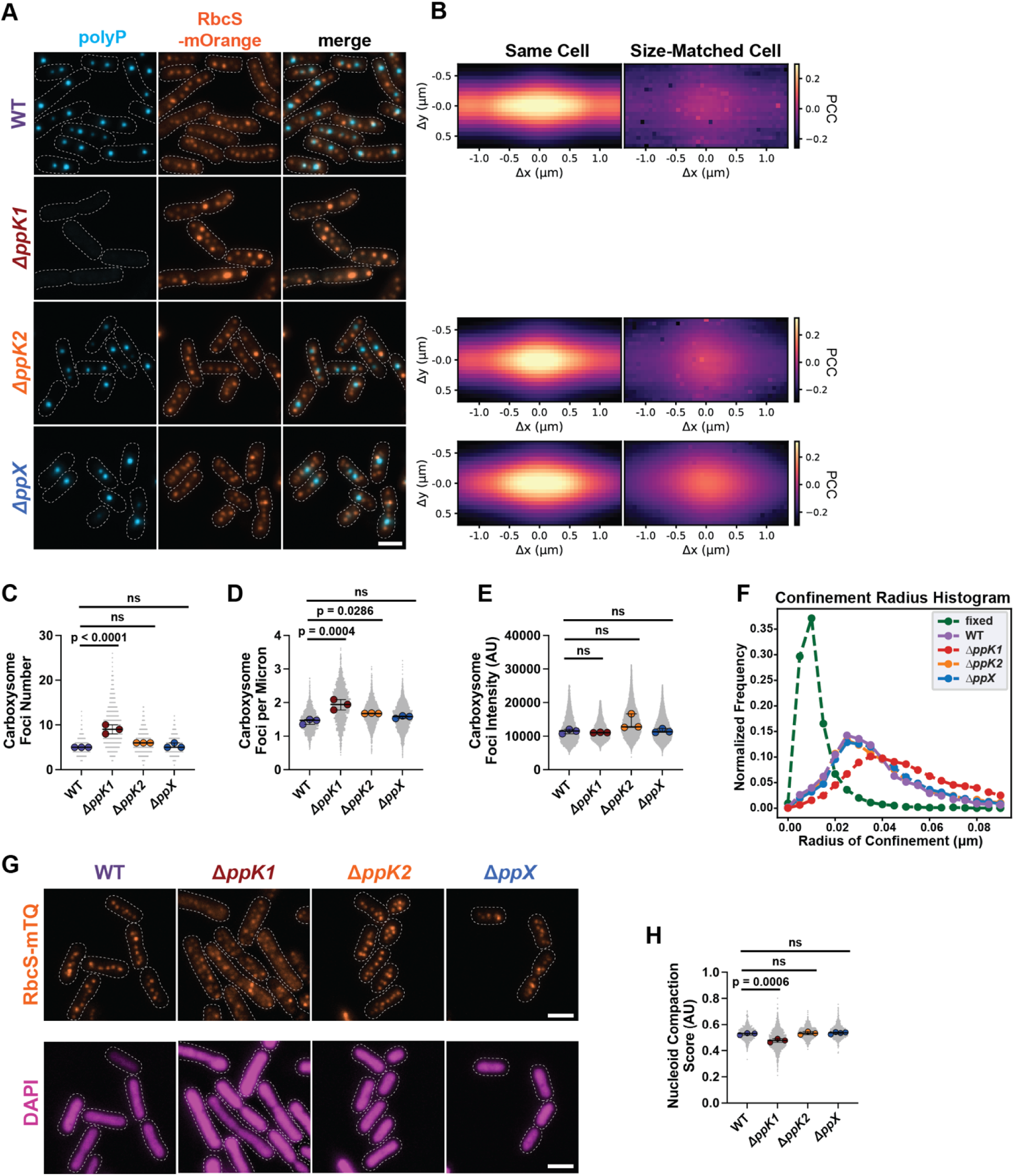
Δ*ppK1* disrupts carboxysome organization, dynamics, and nucleoid compaction. **(A)** Microscopy images of polyP granules (cyan) and carboxysomes (orange), and the merge shows the overlap of the 2 channels for WT, Δ*ppK1*, Δ*ppK2*, and Δ*ppX*. Dotted white line represents the cell boundary from phase contrast (scale bar, 2 μm). WT images are the same as Figure 2A. **(B)** Normalized cross-correlation analysis (see description in Methods section) of carboxysomes and polyP granules from the same cell (top panel) and size-matched cell (bottom panel) for WT, Δ*ppK2*, and Δ*ppX*. Color indicates the Pearson correlation coefficient (PCC) and the x- and y- axes demonstrate shifts of the carboxysome channel. WT panels are duplicates of Figure 2G. **(C)** Number of carboxysomes per cell. Significance from one-way ANOVA and Dunnett’s multiple comparisons. n = 3 biological replicates, bars show median and interquartile range. **(D)** Carboxysomes per unit of cell length, calculated by dividing the number of foci by cell length. Significance from one-way ANOVA and Dunnett’s multiple comparisons. n = 3 biological replicates, bars show median and interquartile range. **(E)** Intensity of carboxysome foci (arbitrary units). Significance from one-way ANOVA and Dunnett’s multiple comparisons. n = 3 biological replicates, bars show median and interquartile range. **(F)** Confinement radius histogram illustrating carboxysome confinement in WT, Δ*ppK1*, Δ*ppK2*, Δ*ppX*, and fixed cells based on TrackMate analysis of fluorescence microscopy videos. **(G)** Microscopy images of carboxysomes (orange) and DAPI-stained nucleoid (magenta) for WT, Δ*ppK1*, Δ*ppK2*, and Δ*ppX*. Dotted white line represents the cell boundary from phase contrast (scale bar, 2 μm). **(H)** Nucleoid Compaction Score (arbitrary units) for WT, Δ*ppK1*, Δ*ppK2*, and Δ*ppX*. Significance from one-way ANOVA and Dunnett’s multiple comparisons. n = 3 biological replicates, bars show median and interquartile range.

Δ*ppK2* and Δ*ppX* cells displayed carboxysome organization similar to WT, with modest increases in carboxysome–polyP proximity **(Fig. 6A-B)**. Δ*ppK1* cells on the other hand, lacked polyP granules and displayed irregularly positioned carboxysomes **(Fig. 6A)**.

Compared to WT and other mutants, Δ*ppK1* cells had significantly more carboxysomes **(Fig. 6C)** with tighter spacing **(Fig. 6D)** and slightly decreased intensity **(Fig. 6E)**. Live-cell imaging followed by trajectory analysis revealed that carboxysomes in Δ*ppK2* and Δ*ppX* cells maintained WT-like dynamics, whereas Δ*ppK1* cells displayed carboxysomes with a larger radius of confinement, indicating enhanced mobility **(Fig. 6F, Video 1)**. Together, the data show that the loss of polyP granules results in more numerous, smaller, and more dynamic carboxysomes.

We previously demonstrated that increased carboxysome mobility can result from altered nucleoid organization^35^. Recent studies further suggest that polyP contributes to nucleoid compaction in diverse bacterial species^7,43^. We therefore hypothesized that the enhanced carboxysome mobility observed in Δ*ppK1* cells might stem from changes in nucleoid structure. Consistent with this model, Δ*ppK1* cells exhibited nucleoid expansion relative to WT **(Fig. 6G-H)**, paralleling the increase in carboxysome mobility **(Fig. 6F)**. Together, these findings indicate that polyP not only directly associates with carboxysomes but also contributes to nucleoid compaction. In doing so, polyP may indirectly regulate carboxysome positioning and dynamics - revealing an unrecognized structural role for polyP in the spatial organization of bacterial carbon-fixation.

### PolyP metabolism influences both the light and dark reactions of photosynthesis

Having established that PPK1-dependent polyP formation regulates nucleoid architecture and carboxysome dynamics, we next asked whether polyP metabolism more broadly shapes photosynthetic cell organization. To assess global ultrastructural consequences, we performed transmission electron microscopy on sectioned cells from each mutant background. While Δ*ppK2* and Δ*ppX* retained carboxysome–polyP associations, Δ*ppK1* cells lacked polyP granules **(Fig. 7A-D)**. Δ*ppK1* cells also exhibited elongated morphology, and consistent with our live-cell microscopy analyses, contained numerous small carboxysomes positioned primarily along the periphery of an expanded nucleoid **(Fig. 7B)**.

**Figure 7.**
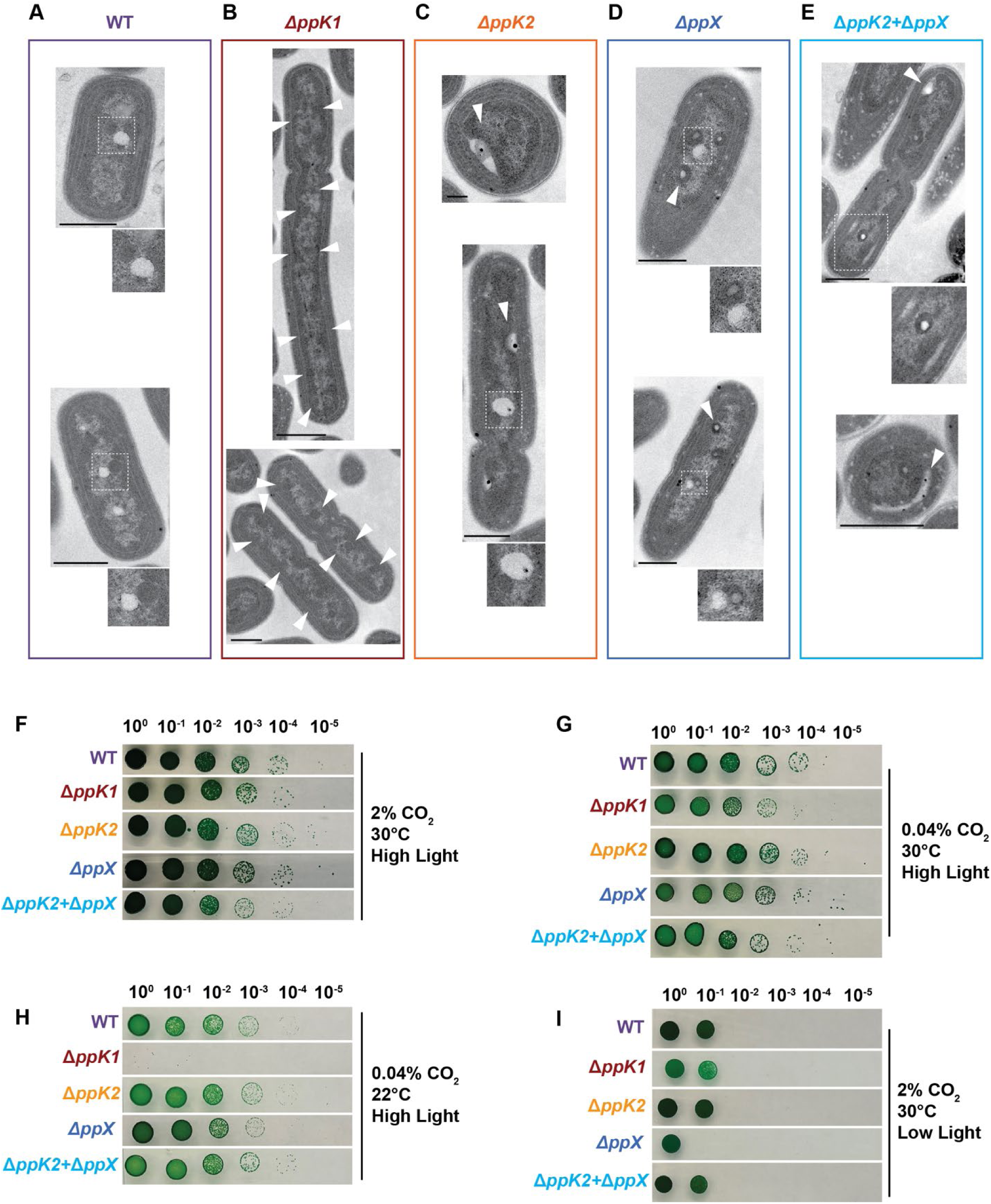
PolyP influences the light and dark reactions of photosynthesis. (A-E) Electron micrographs of **(A)** WT, **(B)** Δ*ppK1* (arrows - carboxysomes), **(C)** Δ*ppK2* (arrows - polyP-like pockets), **(D)** Δ*ppX* (arrows - polyP-like bodies inside of carboxysomes), and **(E)** Δ*ppK2*Δ*ppX* (arrows - polyP-like pockets) (see Methods for preparation). Insets illustrate events of carboxysome-polyP interaction (scale bars, WT - (top) 600 nm, (bottom) 600 nm; Δ*ppK1* - (top) 1 μm, (bottom) 600 nm; Δ*ppK2* - (top) 200 nm, (bottom) 600 nm; Δ*ppX* - (top) 500 nm, (bottom) 500 nm; and Δ*ppK2*Δ*ppX* - (top) 600 nm, (bottom) 1μm). **(F-I)** Series dilution of WT, Δ*ppK1*, Δ*ppK2*, Δ*ppX*, and Δ*ppK2*Δ*ppX* cultures grown in 2% CO2 **(F)**, ambient CO2 **(G)**, ambient CO2 and 22℃ **(H)**, 2% CO2 and 30 µmol m^-1^s^-1^ LED light **(I)**.

Strikingly, additional ultrastructural phenotypes emerged in Δ*ppK2* and Δ*ppX* cells. In Δ*ppK2* cells, polyP-like “pockets” were seemingly wetted to thylakoid membranes **(Fig. 7C, arrows)**. Equally striking, in Δ*ppX* cells, polyP-like inclusions were found within carboxysome lumen **(Fig. 7D, arrows)**. The Δ*ppK2*Δ*ppX* double mutant exhibited both features: polyP granules wetted to thylakoid membranes and polyP-like inclusions within carboxysomes **(Fig. 7E, arrows)**. These observations suggest that polyP metabolism interfaces with both thylakoid architecture (light reactions) and carboxysome structure (dark reactions).

To determine whether these ultrastructural perturbations translate into physiological consequences, we performed serial dilution growth assays at 30°C under high light with either elevated (2%) **(Fig. 7F)** or ambient (0.04%) CO_2_ **(Fig. 7G)**. Under 2% CO_2_ — conditions in which the carbon-concentrating function of carboxysomes is dispensable — all strains grew comparably. However, under ambient CO_2_, Δ*ppK1* exhibited an approximately 100-fold growth defect, demonstrating that polyP is essential for robust growth under physiologically relevant carbon limitation. We next imposed conditions that further constrain photosynthetic efficiency. Upon temperature downshift from 30°C to 22°C under ambient CO_2_, Δ*ppk1* displayed at least a 10^6^-fold growth defect **(Fig. 7H)**, dramatically amplifying the carbon-limitation phenotype and underscoring a heightened requirement for polyP under reduced photosynthetic flux.

Finally, to specifically probe the role of polyP metabolism in the light reactions of photosynthesis, we next imposed low-light stress conditions. Under low-light stress (30 µmol m⁻² s⁻¹) at 2% CO_2_, Δ*ppK1* exhibited only a minor growth defect (<10-fold), whereas Δ*ppX* displayed a 10-fold defect relative to WT **(Fig. 7I)**. Notably, this phenotype was rescued in the Δ*ppK2*Δ*ppX* double mutant, suggesting that PPK2 activity contributes to the growth defect in the absence of PPX. Collectively, these findings indicate that polyP metabolism and its intracellular organization are functionally integrated into photoautotrophic physiology, influencing both thylakoid-associated light reactions and carboxysome-dependent carbon fixation.

## DISCUSSION

Polyphosphate (polyP) is an ancient and highly conserved polymer implicated in diverse aspects of bacterial physiology, including stress adaptation, virulence, and, more recently, nucleoid organization^8^. But how polyP contributes to the spatial and functional architecture of photoautotrophic cyanobacteria remains unclear. Here, we dissect the spatial, structural, and physiological relationships among polyP granules, carboxysomes, and the nucleoid in *S. elongatus* **(Fig 8)**. We show that polyP granules are regularly organized over the nucleoid and frequently associate with carboxysomes that are distributed by McdAB **(Fig 8A)**. Strikingly, carboxysome–polyP interactions are enhanced in the absence of McdAB-mediated positioning **(Fig 8B)**, revealing that active carboxysome positioning normally limits their polyP association. In contrast, polyP localization does not depend on carboxysomes or their spatial distribution.

**Figure 8.**
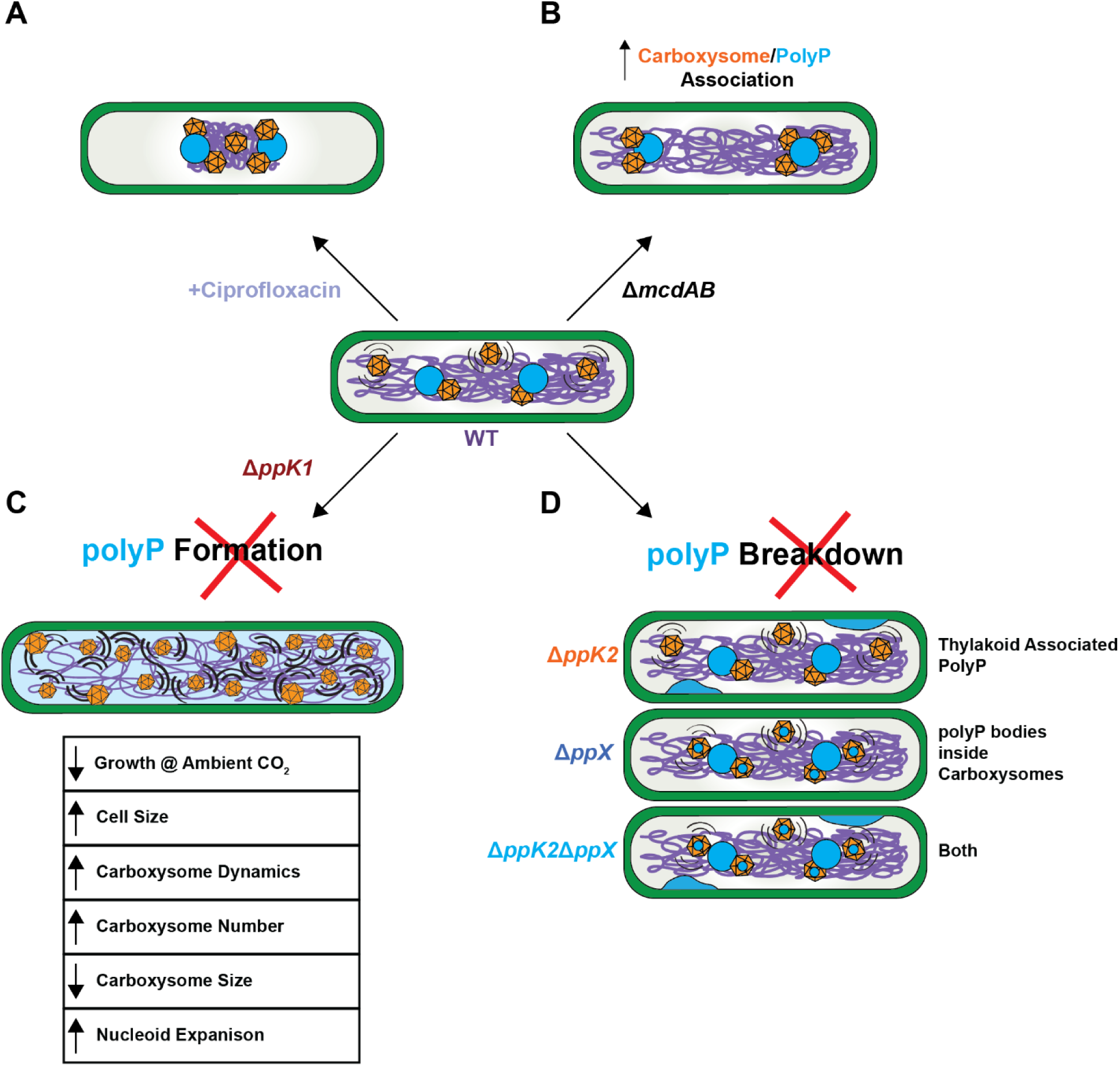
PolyP contributes to carboxysome organization through direct and indirect mechanisms. **(A)** PolyP granules remain nucleoid-associated after ciprofloxacin treatment. **(B)** Disrupting carboxysome positioning increases polyP-carboxysome association. **(C)** Disrupting polyP formation affects carboxysome dynamics and function **(D)** Perturbing polyP breakdown affects the machinery for light and dark reactions of photosynthesis.

Functionally, polyP granules are required for robust growth under ambient CO_2_ conditions and for maintaining proper carboxysome assembly, spatial organization, and nucleoid compaction **(Fig 8C)**. Moreover, perturbing polyP turnover uncovers additional roles in thylakoid and carboxysome architecture **(Fig 8D)**. Collectively, our findings demonstrate that polyP is not merely a storage polymer but a spatially organized regulator of nucleoid compaction, carboxysome organization, and photoautotrophic growth.

### PolyP granules are nucleoid-associated and periodically organized

Through electron microscopy studies, polyP granules have consistently been visualized in close proximity to DNA^14–17,33^, with fiber-like connections in some instances^17^. We demonstrate that polyP granules exhibit regular intracellular positioning in exponentially growing cells, consistently localizing over the nucleoid. Importantly, polyP remains nucleoid-associated even under drug-induced compaction **(Fig 8A)**, arguing against a model in which polyP simply occupies DNA-free cytoplasmic gaps. Instead, our findings support a model in which polyP is directly or indirectly coupled to chromosomal organization. The periodic spacing of polyP granules along the nucleoid resembles positioning patterns mediated by ParA-family ATPases for other intracellular cargos, including plasmids and carboxysomes^44^. McdA is the only ParA-family ATPase encoded by *S. elongatus*, and we show here that polyP organization does not depend on carboxysomes or the McdAB system. These results indicate that polyP localization is governed by a distinct, nucleoid-dependent mechanism independent of ParA-family positioning systems.

The molecular basis underlying polyP spatial organization remains unclear. In *Pseudomonas aeruginosa*, polyP positioning has been linked to the histone-like protein AlgP^23^, yet no AlgP homolog is present in *S. elongatus*. In other well-studied organisms, including *Escherichia coli* and *Caulobacter crescentus*, the mechanism governing polyP localization also remains undefined. Notably, proteomic analyses of isolated polyP granules from *P. aeruginosa* identified proteins containing a <u>c</u>onserved <u>h</u>istidine <u>α</u>-helical <u>d</u>omain (CHAD)^23^. CHAD proteins are widely conserved across bacteria and localize to polyP granules^45,46^. These proteins have been termed “phosins” due to their similarity to phasins, which organize PHB granules; however, their precise function remains unknown, and in several organisms, deletion of phosin-like proteins does not substantially disrupt polyP granule formation^45^. Thus, despite the identification of candidate polyP-associated proteins, the unifying principles that govern polyP organization across bacteria remain unresolved.

Additionally, polyP has been implicated in cell cycle regulation, raising the possibility that this periodic patterning is coordinated with chromosome dynamics or cell division machinery^5,12^. PolyP has also been shown to interact with nucleoid-associated proteins (NAPs), providing a potential molecular link between polyP and DNA^7^. Given the highly negative charge of both polyP and DNA, metal cofactors likely facilitate or stabilize these interactions^48^. Intriguingly, we find that loss of polyP granules leads to nucleoid expansion, suggesting a more direct structural role whereby polyP itself contributes to nucleoid compaction. Similar nucleoid expansion phenotypes following disruption of polyphosphate kinases have been observed in other bacteria^5,43^, indicating that this relationship may be broadly conserved. Alternatively, the absence of polyP could indirectly alter nucleoid structure by affecting NAP availability or DNA-binding dynamics^7^. Together, these observations point to polyP as a modulator of chromosome architecture, though the precise molecular mechanisms remain to be defined.

### PolyP and carboxysome positioning are coordinated but mechanistically independent

Although prior ultrastructural studies reported proximity between polyP granules and carboxysomes^16,18^, whether this association was functional or incidental remained unclear.

Using live-cell imaging and quantitative cross-correlation analysis, we demonstrate that polyP and carboxysomes frequently associate. Despite this association, we find that polyP positioning does not depend on carboxysomes or McdAB-mediated carboxysome distribution. These results rule out a hitchhiking model in which polyP relies on carboxysome positioning machinery.

Interestingly, the reciprocal relationship does not hold. In the absence of McdAB, carboxysomes cluster more frequently around polyP granules, and TEM reveals multiple carboxysomes aggregated around single polyP bodies. This suggests that McdAB positioning actively counterbalances an intrinsic affinity between carboxysomes and polyP. Thus, beyond distributing carboxysomes across the nucleoid, McdAB appears to function as an anti-aggregation system^49^, preventing excessive carboxysome coalescence at polyP granules.

Across multiple organisms, polyP granules localize preferentially to the quarter positions of the cell^5,12^, suggesting a conserved spatial logic. Notably, carboxysome biogenesis in *S. elongatus* has also been observed at these quarter positions^47^, raising the possibility of a functional link between the two. One attractive model is that polyP granules act as nucleation hubs for *de novo* carboxysome assembly, after which the McdAB system separates and distributes newly formed carboxysomes across the nucleoid.

The physical nature of the polyP-carboxysome association remains unresolved. Our study shows that it is not McdA or McdB, because in the absence of these proteins, carboxysomes cluster more frequently with polyP. Prior work in *H. neapolitanus* described a lattice-like linker between these structures^18^. Candidates in *S. elongatus* may include shell proteins, CHAD proteins^45,46^, or even PPK1 itself, which forms filamentous structures *in vitro*^50,51^. Future proteomic and ultrastructural studies will be required to identify the molecular connector.

### PolyP regulates nucleoid architecture and carboxysome dynamics

Deletion of *ppk1* eliminates polyP granules, resulting in expanded nucleoids and an increased number of smaller carboxysomes with altered spacing and enhanced mobility. These phenotypes parallel our previous observations linking nucleoid compaction state to carboxysome organization^35^. Together with emerging evidence that polyP modulates chromosome structure in diverse bacteria^7,43^, our findings support a model in which polyP contributes nucleoid architecture. In this framework, polyP influences carboxysomes through two interconnected mechanisms: (1) Direct association — where polyP granules physically contact carboxysomes, and (2) Indirect regulation — where polyP-mediated nucleoid compaction constrains carboxysome positioning and mobility governed by the McdAB system. Loss of polyP destabilizes nucleoid organization, leading to increased carboxysome mobility and altered spatial distribution.

### PolyP metabolism interfaces with both light and dark reactions of photosynthesis

We find several interesting structural phenotypes in cells that do not have polyP or are unable to break down granules, which shed light on the potential metabolic need for carboxysome-polyP interactions. Δ*ppK1* cells, which we show lack polyP granules, possess more carboxysomes that are smaller. This suggests that a compacted nucleoid may be important for carboxysome biogenesis, or polyP granules themselves may act as a nucleation hub for the proper assembly of carboxysomes. Consistent with this proposal, *in vitro* assembly studies using purified hexameric shell proteins required phosphate-containing buffers and the authors suggested that incorporation of phosphate groups between hexamer units is critical for proper shell tiling^52^. This supports a potential role for phosphate in carboxysome biogenesis by facilitating shell assembly.

Interestingly, in Δ*ppX* cells, we observed the presence of polyP-like bodies inside of carboxysomes, further pointing to a role for polyP during the inside-out assembly mechanism of carboxysomes during their biogenesis in β-cyanobacteria^42^. PolyP-like bodies have been previously observed inside of carboxysomes in WT *S. elongatus* cells under low light growth conditions, highlighting a potential functional role during photosynthetic stress^53^. Additionally, for Δ*ppK2* cells, we found polyP pockets that localized along the thylakoid membrane, alluding to a potential role for polyP in the light reaction of photosynthesis, specifically the photosystem machinery.

Functionally, polyP is dispensable under elevated CO_2_ but essential under ambient CO_2_, where Δ*ppK1* cells exhibit severe growth defects that are dramatically amplified at lower temperatures. These conditions constrain photosynthetic flux, highlighting a critical role for polyP under physiologically relevant carbon limitation. Other groups have also shown that Δ*ppK* cyanobacterial species have reduced fitness compared to WT under non-supplemented CO_2_ conditions^50,56^. PolyP depletion may alter cellular energy states; cells lacking polyP have been reported to exhibit elevated ATP levels and a more dynamic cytoplasm^43^. PolyP may also buffer cytoplasmic pH or influence phosphate availability required for Rubisco activation^18,54,55^, providing additional biochemical links between polyP metabolism and carboxysome function. Additionally, in *ΔppK* cells grown under CO_2_-supplemented conditions, several carboxysome shell proteins were upregulated, raising the possibility that carboxysomes do not assemble properly in this background^56^. This observation further supports our findings and proposal that polyP plays an important role in carboxysome biogenesis.

Under low-light stress, Δ*ppX* and, to a lesser extent Δ*ppK1*, exhibits growth defects. The Δ*ppX* growth defect was rescued by deletion of *ppk2*, suggesting that polyP turnover dynamics, rather than simply polyP abundance, influence light-reaction fitness. PolyP has also been linked to stringent response pathways^6,11,32,60,61^, and basal ppGpp regulation is necessary for proper growth and metabolism in *S. elongatus*^62^, suggesting that polyP could help to maintain a constant low-level stress metabolism in circadian-controlled organisms and further connecting phosphate metabolism to stress and energy homeostasis. Collectively, these findings indicate that polyP metabolism integrates phosphate storage, nucleoid structure, and photosynthetic machinery across both light and dark reactions.

### PolyP as an architectural and metabolic integrator

We propose a model in which polyP granules function as multifunctional nodes that integrate chromosome organization, carboxysome positioning, and photosynthetic metabolism. In WT cells, polyP localizes along the nucleoid, contributes to nucleoid compaction, and associates transiently with carboxysomes. McdAB positioning distributes carboxysomes across this scaffold, preventing aggregation at polyP. In the absence of polyP, nucleoid expansion leads to compromised carboxysome organization and carbon fixation function under ambient CO_2_ conditions. In the absence of McdAB, carboxysomes collapse onto polyP granules, revealing a latent affinity normally restrained by active positioning. Together, our findings redefine polyP granules as spatially organized regulators of bacterial cell architecture rather than passive storage inclusions. By linking phosphate metabolism to nucleoid structure and carbon fixation machinery, polyP emerges as a central organizing principle of the photoautotrophic cytoplasm.

### Conclusions and future directions

This work raises several key questions. What molecular factors mediate polyP positioning in bacteria? What is the physical basis of polyP-carboxysome association? Does polyP directly regulate Rubisco activation, pH buffering, or energy flux within carboxysomes? How does polyP influence carboxysome biogenesis? Elucidating these mechanisms will uncover fundamental principles governing the coordination of metabolic polymers and proteinaceous organelles in bacterial cells. More broadly, our findings suggest that metabolic polymers such as polyP may act as architectural regulators of intracellular organization, extending beyond their classical roles in storage and stress response. Understanding this interplay will deepen our insight into bacterial subcellular organization and the integration of metabolism with spatial control.

## METHODS AND MATERIALS

### Growth conditions

All cultures were grown in BG11 media pH 8.3 (Sigma), with regular back-dilutions into fresh BG11 media, to maintain exponential growth. Cultures were maintained at a 50 mL volume in 125 mL beveled flasks (Corning), or 100 mL in 250 mL beveled flasks. A Minitron incubator (Infors-HT) was used to grow the cultures at 30℃, 2% CO_2_, and 130 RPM shaking, with 60 µmol m^-1^s^-1^ LED light.

### Spot Serial Dilutions

Exponentially growing cultures were back diluted to an OD_750_ of ∼0.2. Using PCR strip tubes, the cultures were diluted sequentially in 180 µL BG11 media. Using a multi-channel pipette, dilutions were spotted at a volume of 5 µL on a BG11 agar plate. Plates were then transferred to an incubator at 30℃, 60 µmol m^-1^s^-1^ LED light, with or without supplemented 2% CO_2_, 22℃, 60 µmol m^-1^s^-1^ LED light, without supplemented 2% CO_2_, or 30℃, 30 µmol m^-1^s^-1^ LED light, with supplemented 2% CO_2_. Plates were grown until no additional colonies grew.

### DAPI staining

To visualize the nucleoid, 2 mL of cells were harvested by centrifugation at 16,000 x g for 1 min. After washing cells once with 1X PBS (pH 7.2), cells were resuspended in 100 µL PBS. DAPI was added to achieve a final staining concentration of 9 µg/mL. Cells were then incubated in the dark for 20 minutes at 30℃, 2% CO_2_, and 130 RPM shaking. Prior to visualization, DAPI-stained cells were washed twice with 1 mL H_2_O and resuspended in 100 µL H_2_O.

For polyphosphate visualization, DAPI was added to 2 mL of culture at a final staining concentration of 10 µg/mL. DAPI-stained cells were then incubated in the dark for 4 hours at 30 ℃, 2% CO_2_, and 130 RPM shaking. Cells were then rinsed twice with 1 mL H_2_O before plating for imaging.

### Live fluorescence microscopy

To image, ∼1 µL of cells was spotted on an agarose disk composed of 1.5% UltraPure agarose (Invitrogen) in BG11. The disk was then transferred to a 35 mm cover glass-bottom dish (MatTek). Images were captured using the NIS Elements software on a Nikon Ti2-E inverted microscope with a 100x objective lens, SOLA LED light source, and a Hamamatsu camera.

Carboxysomes were captured using a CFP filter, for RbcS-mTQ, or mOrange filter, for RbcS-mOrange, DAPI stained DNA with a DAPI filter, and DAPI stained polyP with a CFP filter.

### Transmission electron microscopy sample preparation without fixation

For whole cell mounting without fixation, strains were grown to exponential phase. 5 µL of unfixed culture was then deposited onto a formvar coated carbon grid. The culture was incubated on the grid for 2.5 - 5 min before wicking away the excess using Kimwipes. The grid was then washed 5X with 5 µL deionized water. Water was immediately wicked away between washes using a Kimwipe. The grid then was allowed to dry for ∼ 2.5 min, ensuring that it was visibly dry before storing. Grids were prepared from 2 biological replicates for each strain. This protocol was adapted from Chawla et al. 2022.

### Transmission electron microscopy sample preparation with fixation

Strains were grown to exponential phase and 6 mL of culture was collected by centrifugation at 16,000 x g for 1 min. Cells were then rinsed with 1 mL 1X PBS pH 7.4 and centrifuged at 16,000 x g for 1 min. After removing the supernatant, the pellet was resuspended in 2 mL of fixative (2.5% glutaraldehyde, 2% formalin in 0.1M sodium cacodylate buffer (CB)). Samples were then fixed for 2 hours at room temperature before being stored at 4℃ and then delivered to the University of Michigan Microscopy Core. Samples were then embedded in 4% agarose, washed with 0.1M CB for 15 min 3 times, incubated in 2% OsO4 in 0.1M CB for 1 hour, washed with 0.1M CB for 15 min 3 times, washed with 0.1M acetate buffer (AB) for 5 minutes 3 times, incubated with 2% Uranyl Acetate (UA) in 0.1M AB for 1hour, washed with 0.1M AB, for 5 minutes x2, and rinsed with deionized water for 5 min. A dehydration series was then performed where the samples were incubated for 15 min at 4°C in varying ethanol solutions: 30, 50, 70, 80, 90, 95, 100, 100, Acetone. Samples were then incubated in acetone:resin solution at a 2:1 ratio for 1 hour, a 1:1 ratio for 2 hours, and a 1:2 ratio for 16 hours. They were then kept in 100% resin under vacuum conditions for 48 hours. Samples were then embedded and polymerized in a 70°C oven for 24 hours. The resulting blocks were then sectioned at a thickness of 70 nm.

### Transmission electron microscopy

Grids were imaged on a Jeol 1400+ Electron microscope. For sectioned TEM samples, grids were imaged at 60 kV. For whole cell non-fixed samples, grids were imaged at 120 kV to visualize polyP granules^23^.

### Image Analysis

#### TrackMate

Using the FIJI plugin TrackMate, carboxysomes were detected and tracked during videos. The LoG detector was used with an estimated blob diameter of 3 pixels and a threshold of 80.0. The Simple LAP tracker was used with settings of 7.5 pixels for Max Distance, 4.5 pixels for Gap-Closing Max Distance, and 2 frames for the Gap-Closing Max Frame Gap. For downstream carboxysome analysis, the Spots in Track Statistics was primarily used.

#### Confinement radius

As described previously, the first ten xy localizations per carboxysome track were used to calculate the confinement radius by finding the average xy position across these locations^35^. The confinement radius is the average difference between the calculated xy position and the track coordinates.

#### Cell registration

Before analyzing videos in TrackMate, cell registration was completed to correct for any drift that occurred during the video. The chlorophyll channel was registered to the first frame using shifts measured by cross-correlation. The shift for each frame was then applied to the corresponding frame in the carboxysome channel. A registered stack was produced and saved as a separate TIFF file that was then used for downstream analysis.

#### Cell segmentation

Phase-contrast images were used for cell segmentation in the Omnipose package in Python. “bact_phase” was used as the pre-trained model with detection settings of -3.74 for the mask threshold and 0.00 for the flow threshold. Using the Omnipose GUI, false cell masks were removed, and additional masks were added manually.

#### Fluorescent focus detection

Carboxysomes and polyphosphate granules were detected using a custom Python Script as described previously^35^. The preprocessing settings can be found in **Supplementary Table 1**.

#### Nucleoid morphology

Nucleoid compaction was calculated as described previously^35^. Nucleoid compaction score *s*_comp_ was calculated by subtracting the ratio of the nucleoid area to cell area from 1: *s*_comp_ = 1 – (*A*_nuc_/*A*_cell_).

#### Normalized cross-correlation analysis

The degree of association between carboxysomes and polyphosphate granules was quantified by applying normalized cross-correlation (NCC) analysis to images of labeled carboxysomes and polyphosphate granules^63^. The normalized cross-correlation function, *R*, is a measurement of the correlation between two images, *I_cbs_* and *I_ppg_*, shifted relative to each other by *Δx* pixels in *x* and *Δy* pixels in *y*:

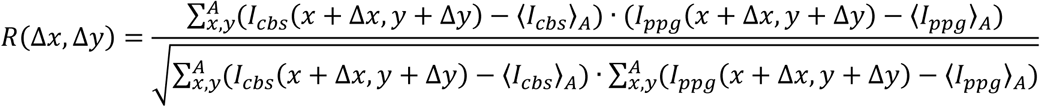

*A* is the set of pixels where the segmented region of interest overlaps with itself, shifted by *Δx* and *Δy*.

We calculated the NCC for each individual cell. Prior to NCC calculation, cell images were rotated so that the long axis of the cell was aligned with the x-axis. This convention makes it so that the shape of the NCC in *Δx* informs on the spacing between foci along the long axis of the cell, and the shape in *Δy* informs on the spacing between foci along the short axis. Finally, NCCs were averaged across all cells for each condition.

#### Carboxysome and polyphosphate heatmaps

The localization coordinates of detected carboxysome and polyphosphate granule foci and cell masks were used to make localization density heatmaps with the spideymaps tool (https://github.com/BiteenMatlab/spideymaps) in Python as described previously^35^. Heatmaps were normalized by dividing by the average density over the entire map so that the displayed map values represent the density relative to a perfectly uniform distribution of particles. To visualize the dependence of foci positioning on cell length, cells were grouped according to their lengths prior to the calculation of the heatmaps.

## ACKNOWLEDGEMENTS

This work was supported by the National Institute of General Medical Sciences of the National Institutes of Health to J.S.B. (Award # R01-GM144731) and A.G.V. (Award # R35-GM152128). D.F.S. is an Investigator in the Howard Hughes Medical Institute. C.E.D. acknowledges funding from the Rackham Candidate Grant.

Conceptualization, C.E.D., D.F.S., and A.G.V.; Investigation, C.E.D.; Software, D.J.F.; Supervision, J.S.B. and A.G.V.; Writing – original draft, C.E.D. and A.G.V.; Writing – review & editing, C.E.D., D.J.F., D.F.S., J.S.B., A.G.V.

