## Supplementary Information for "Polyphosphate acts as an architectural regulator of carbon fixation and nucleoid structure in cyanobacteria"

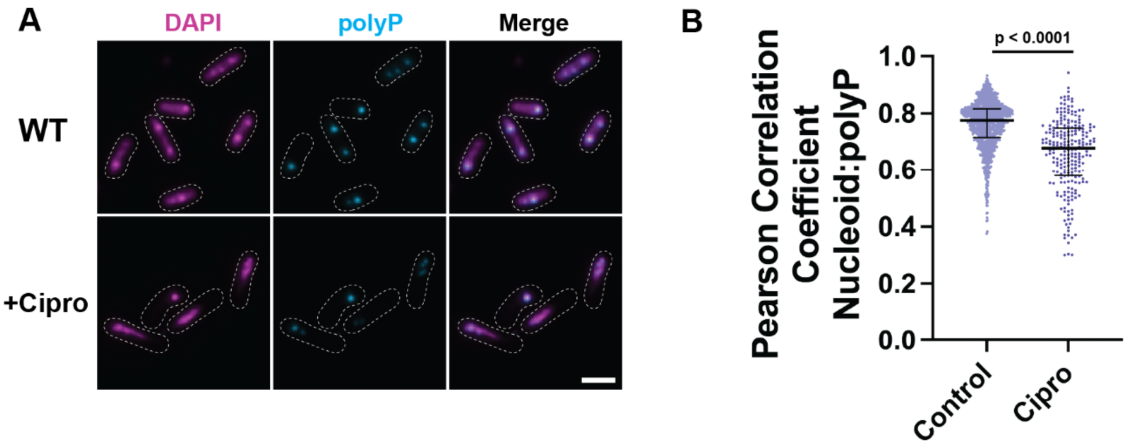


**Figure S1 - PolyP granules remain nucleoid-associated after ciprofloxacin treatment. (A)** Microscopy images of WT exponential cells without (top row) and with ciprofloxacin treatment (bottom row), DAPI-stained nucleoid (magenta), polyP (cyan), and merge is the overlap of the 2 channels. Dotted white line represents the cell boundary from phase contrast (scale bar, 2 μm). **(B)** Pearson correlation coefficient of DAPI-stained nucleoid channel and polyP channel. Significance from Mann-Whitney. n = 3 biological replicates, bars show median and interquartile range.

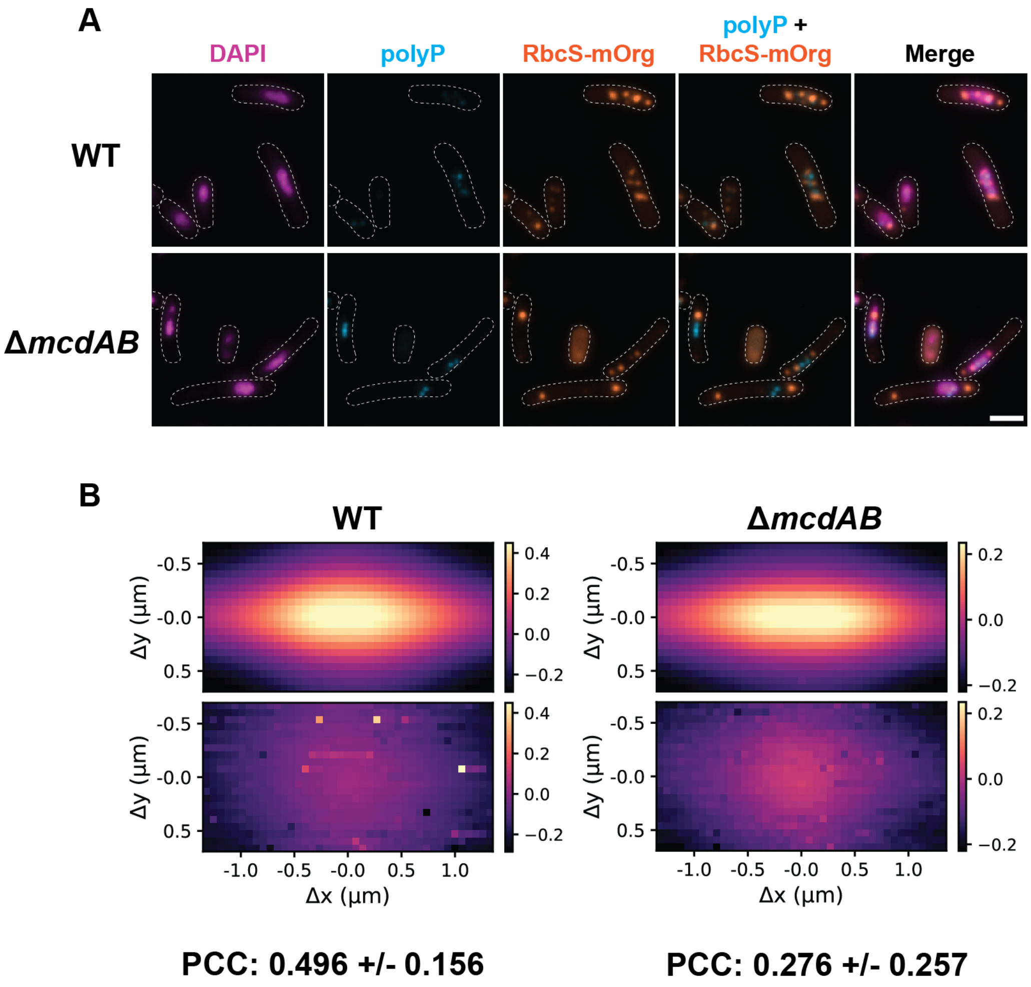


**Figure S2 - Nucleoid compaction in Δ*mcdAB* cells can reduce carboxysome-polyP association. (A)** Microscopy images of WT exponential cells (top row) and Δ*mcdAB* cells (bottom row) after ciprofloxacin treatment. DAPI-stained nucleoid (magenta), polyP (cyan), carboxysomes (orange), overlay of polyP and carboxysome channels (cyan and orange) and merge is the overlap of all 3 channels. Dotted white line represents the cell boundary from phase contrast (scale bar, 2 µm). **(B)** Normalized Cross-Correlation Analysis (see description in Methods section) of carboxysomes and polyP granules from the same cell (top panel) and size-matched cell (bottom panel) for WT cells (left) and Δ*mcdAB* cells (right) after ciprofloxacin treatment. Color indicates the Pearson correlation coefficient (PCC), and the x- and y- axes demonstrate shifts of the carboxysome channel. The average PCC is reported below with the standard deviation.

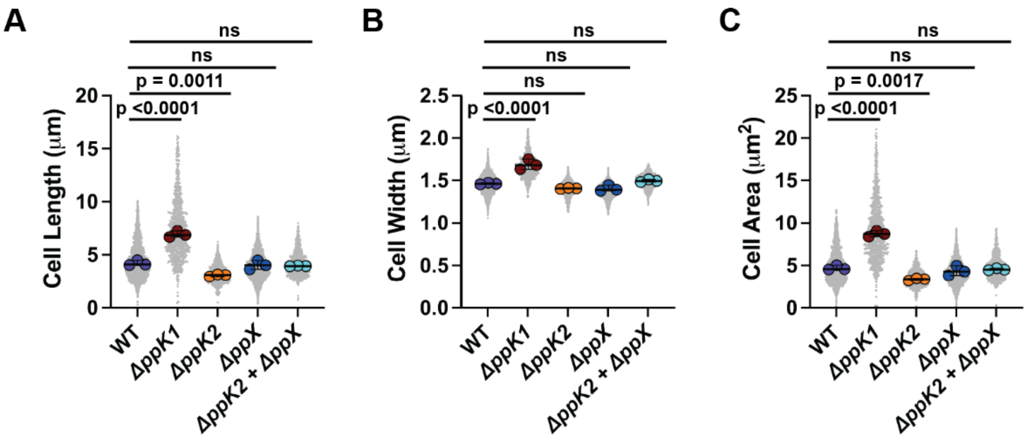


**Figure S3 - Δ*ppK1* cells display altered cell morphology. (A)** Cell lengths for WT, Δ*ppK1*, Δ*ppK2*, Δ*ppX*, and Δ*ppK2*Δ*ppX* cells. **(B)** Cell widths for WT, Δ*ppK1*, Δ*ppK2*, Δ*ppX*, and Δ*ppK2*Δ*ppX* cells. **(C)** Cell areas for WT, Δ*ppK1*, Δ*ppK2*, Δ*ppX*, and Δ*ppK2*Δ*ppX* cells. All plots display significance from one-way ANOVA and Dunnett's multiple comparisons. n = 3 biological replicates, bars show median and interquartile range.

### VIDEO LEGENDS

**Video 1. Carboxysome dynamics in polyP metabolism mutants of *S. elongatus***. Phase contrast channel overlaid with carboxysomes (RbcS-mTQ, LUT is orange) in the following strains - WT (far left), ∆*ppK1* (center left), ∆*ppK2* (center right), ∆*ppX* (far right).One-hour video at 1 frame every 3 minutes. Playback speed is 10 frames per second (scale bar is 2 μm).

### SUPPLEMENTARY TABLES

**Supplementary Table 1 -** Python Analysis Settings

| **Focus Detected** | **Background Subtraction** | **Image Denoise** | **Add Gaussian Blur** | **Detection** |
|---|---|---|---|---|
| RbcS-mOrg Carboxysomes | restoration.rolling_ball(cell_fluor,radius = 3) | filters.unsharp_mask(cell_fluor_bg, radius = 2, amount = 5) | filter.gaussian(cell_fluor_sharp, 0.01) | min_sigma = 2.75, max_sigma = 3, threshold = 0.005, overlap = 0.75 |
| RbcS-mTQ Carboxysomes | restoration.rolling_ball(cell_fluor,radius = 3) | filters.unsharp_mask(cell_fluor_bg, radius = 2, amount = 5) | filter.gaussian(cell_fluor_sharp, 0.01) | min_sigma = 1.75, max_sigma = 3, threshold = 0.0025, overlap = 0.75 |
| polyP | restoration.rolling_ball(cell_fluor,radius = 3) | filters.unsharp_mask(cell_fluor_bg, radius = 3, amount = 5) | filters.gaussian(cell_fluor_sharp, .5) | min_sigma = 2.5, max_sigma = 5, threshold = 0.0125, overlap = 0.95 |

**Supplementary Table 2 -** Constructs used

| **Strain** | **Identifier** | **Description** | **Reference** |
|---|---|---|---|
| WT | C1 | WT *Synechococcus elongatus* PCC 7942 | |
| Δ*ccm* | C103 | mNG-McdB + RbcS-mTQ + ΔccmK2-O | [20] |
| Δ*mcdA* | C7 | Native deletion of mcdA by replacing the gene with a kanamycin resistance cassette | [20] |
| Δ*mcdB* | C8 | Native deletion of mcdB by replacing the gene with a kanamycin resistance cassette | [20] |
| Δ*mcdAB* | C9 | Native deletion of mcdAB by replacing the operon with a kanamycin resistance cassette | [20] |
| RbcS-mTQ | C3 | RbcS-mTQ with the native promoter at neutral site I | [20] |
| RbcS-mOrange2 | C30 | RbcS-mOrange2 with the native promoter at neutral site I | [20] |
| Δ*mcdA* + RbcS-mOrange2 | C107/CED28 | Native deletion of mcdA by replacing the gene with a kanamycin resistance cassette, RbcS-mOrange2 with the native promoter at neutral site I | This paper |
| Δ*mcdB* + RbcS-mOrange2 | C108/CED29 | Native deletion of mcdB by replacing the gene with a kanamycin resistance cassette, RbcS-mOrange2 with the native promoter at neutral site I | This paper |
| Δ*mcdAB* + RbcS-mOrange2 | C109/CED30 | Native deletion of mcdAB by replacing the operon with a kanamycin resistance cassette, RbcS-mOrange2 with the native promoter at neutral site I | This paper |
| Δ*ppK1* | C66/CED11 | Native deletion of ppK1 by replacing the gene with a spectinomycin resistance cassette | This paper |
| Δ*ppK1* + RbcS-mTQ | C69/CED14 | Native deletion of ppK1 by replacing the gene with a spectinomycin | This paper |

| | | | |
|---|---|---|---|
| | | resistance cassette, RbcS-mTQ with the native promoter at neutral site I | |
| Δ*ppK1* + RbcS-mOrange2 | C110/CED31 | Native deletion of ppK1 by replacing the gene with a spectinomycin resistance cassette, RbcS-mOrg with the native promoter at neutral site I | This paper |
| Δ*ppK2* | C122/CED42 | Native deletion of ppK2 by replacing the gene with a kanamycin resistance cassette | This paper |
| Δ*ppK2* + RbcS-mTQ | C125/CED46 | Native deletion of ppK2 by replacing the gene with a kanamycin resistance cassette, RbcS-mTQ with the native promoter at neutral site I | This paper |
| Δ*ppK2* + RbcS-mOrange2 | C127/CED48 | Native deletion of ppK2 by replacing the gene with a kanamycin resistance cassette, RbcS-mOrg with the native promoter at neutral site I | This paper |
| Δ*ppX* | C73/CED18 | Native deletion of ppX by replacing the gene with a spectinomycin resistance cassette | This paper |
| Δ*ppX* + RbcS-mTQ | C76/CED21 | Native deletion of ppX by replacing the gene with a spectinomycin resistance cassette, RbcS-mTQ with the native promoter at neutral site I | This paper |
| Δ*ppX* + RbcS-mOrange2 | C111/CED32 | Native deletion of ppX by replacing the gene with a spectinomycin resistance cassette, RbcS-mOrg with the native promoter at neutral site I | This paper |
| Δ*ppX* + Δ*ppK2* | C124/CED45 | Native deletion of ppX by replacing the gene with a spectinomycin resistance cassette, native deletion of ppK2 by replacing the gene with a kanamycin resistance cassette | This paper |
